# Hydroperiod buffers water surface decline in dryland wetlands: A 36-year analysis in Hwange National Park

**DOI:** 10.64898/2026.04.13.718152

**Authors:** Alexis Roy, Aiten Alava Baldazo, Florence D. Hulot, Kamel Soudani

## Abstract

Drylands are experiencing increasingly intense and frequent drought events due to climate change. Wetlands in drylands are therefore under increasing pressure as their inundation regimes are altered. In southern African savannas, wetlands are often the only sources of free water for the ecosystem. Changes in the hydroperiod may alter vegetation and water surfaces, which could be early signals of desertification in their immediate vicinity. To investigate trends in surface cover around wetlands, we applied linear multispectral unmixing to Landsat pixels located near wetlands in Hwange National Park. We assessed spatial gradients in vegetation, water, and bare soil dynamics from 1986 to 2022. The studied wetlands were also grouped by hydroperiod to test whether the response of each surface cover differed with the reliability of the water resource. Our results show a significant decrease in the water fraction of wetlands with short hydroperiods, which was significantly negatively correlated with increasing temperature. Furthermore, water fraction was significantly positively correlated with vegetation fraction. This correlation suggests that vegetation could be affected if water surfaces continue to decline. Finally, this study is the first to demonstrate a decline in water surfaces in Hwange National Park, with potential implications for wildlife conservation.

## 1. Introduction

The classical ecological framework delineates freshwater and terrestrial ecology as two distinct disciplines. Wetlands, by falling into both categories without completely belonging to both, are often defined as ecotones between freshwater and terrestrial ecosystems (Mitsch and Gosselink, 2007). This definition is based on the presence of standing water at specific periods, associated with unique vegetation adapted to waterlogged soils (Boix et al., 2020). Their dynamic aspect shapes a unique combination of terrestrial and aquatic ecosystem attributes (Mitsch and Gosselink, 2007), providing specific ecosystem services.

Indeed, wetlands are biodiversity hotspots, providing habitats for a wide range of plant and animal species (Sievers et al., 2018; Zedler and Kercher, 2005). The capacity of wetlands to host such species relies on their hydrological dynamics, as they shape the landscape and the species capable of withstanding hydrological variations as well as the available ecological niches (Rolls et al., 2018; Yang et al., 2025; Zhang et al., 2023). Although often overlooked, drylands comprise 45% of the total land surface, spanning from boreal tundra to hyper-arid deserts (Prăvălie, 2016). They are defined as climatic zones with a ratio of annual precipitation to potential evapotranspiration below 0.65, resulting in a potential water deficit (Grenfell et al., 2022; Middleton et al., 1997). Therefore, in drylands, hydrological dynamics in wetlands are more intense, with periods of intense pressure on water resources. These specific climatic conditions led to the definition of a specific type of wetland: Wetlands in Drylands (WiDs) (Furlan et al., 2021; Grenfell et al., 2022; Parra et al., 2021; Tooth and McCarthy, 2007; Williams, 1999). In particular, they were clearly defined by Tooth and McCarthy (2007) as characterized by more frequent and/or prolonged periods of desiccation, higher levels of sedimentation, more frequent fires, and longer developmental histories, sometimes dating back to the Pleistocene. Even if the spatial expansion of drylands due to climate change is disputed (Berg and McColl, 2021; Feng and Fu, 2013; Huang et al., 2016a; Zhang et al., 2024), they experience increasing intensity and frequency of drought events caused by a decrease in precipitation and an increase of the potential evapotranspiration (Huang et al., 2016b, 2016a; IPCC, 2022; Prăvălie, 2016). The increasing aridity may alter WiD’s water dynamics and substantially impact species that rely on specific hydroperiods (Grenfell et al., 2022; Parra et al., 2021), while also modifying the landscapes in which they are integrated.

In regions with limited water availability, vegetation is significantly influenced by the concentration of fauna around WiDs. This interaction creates a vegetation gradient known as the piosphere (Chamaillé-Jammes et al., 2009; Lange, 1969; Thrash and Derry, 1999). While the relationship between piosphere size and water availability has been well studied, often through metrics such as the Normalized Difference Vegetation Index (NDVI) or tree density, this perspective often overlooks crucial dynamics. Specifically, the macroscopic approach may fail to capture the interplay among various surface covers, leading to an incomplete understanding of the proportions of bare soil, vegetation, and water in the piosphere. This represents a significant scientific gap, given that the evolution of the spatial organization of vegetation, the size of patches, and overall vegetation coverage are key indicators of desertification processes, particularly in conjunction with rising temperatures (Barbier et al., 2006; Huang et al., 2021; Maestre and Escudero, 2009). Additionally, the risk of desertification in WiDs is compounded by increased extreme water level fluctuations in the eulittoral zone (Parra et al., 2021). Therefore, it is essential to describe these dynamics and their interactions to better predict potential desertification processes in relation to temperature evolution.

Southern Africa faces one of the highest rates of species extinction (Ceballos et al., 2017), and water availability is a critical conservation concern. This is especially true in Hwange National Park (HNP) in Zimbabwe, where, during the dry season, water sources attract higher concentrations of animals seeking water and prey (Valeix, 2011). Larger mammals impact the vegetation in wetlands through trampling, foraging, nutrient inputs, and uprooting. This leads to the creation of specific vegetation gradients around water sources, called the piosphere (Chamaillé-Jammes et al., 2009; Hulot et al., 2019). Water availability in the park primarily comes from temporary to permanent water points, with high variability of morphological features (Roy, 2025; Roy et al., 2025). Because of its protected status, HNP remains largely untouched by human activity apart from water pumping in some wetlands; however, temperatures are rising significantly, and drought intensity is increasing (Chamaillé-Jammes et al., 2007a; Roy et al., 2025). While the piosphere has been extensively studied in the park, the dynamics of surface cover have not yet been explored (Chamaillé-Jammes et al., 2009; Mpakairi, 2019), particularly in the closed vicinity of water. These factors make HNP an ideal case study for assessing changes in vegetation, bare soil, and water surfaces, particularly in relation to the frequency of water availability, which we anticipate will play a significant role in mitigating the impacts of climate change.

Estimating vegetation, water, and bare-soil dynamics can be done through extensive fieldwork. This approach, however, entails a trade-off with spatial representativeness, as it limits the number of sampled WiDs and, consequently, their overall representativeness. This issue is particularly challenging in remote areas, where monitoring costs are high (Chamaillé-Jammes et al., 2007b). To address this, remote sensing techniques are increasingly used to monitor WiDs in inaccessible locations, effectively detecting patterns of water availability (Roy et al., 2025; Schaffer-Smith et al., 2022). One significant limitation in assessing surface cover in WiDs is the data’s spatial resolution, which may be too coarse to capture the dynamics of these ecosystems. Landsat has been extensively utilized for WiD studies due to its consistent data acquisition since the 1980s (Gxokwe and Mazvimavi, 2023). Recent studies have also leveraged Sentinel-1 and Sentinel-2 data to map and characterize wetlands. Notably, single-date wetland assessments using the Sentinel-2 constellation have shown its potential with high spatial resolutions ranging from 10 to 60 meters (Slagter et al., 2020). However, the spatial and temporal resolutions of multi-spectral sensors may not always be sufficient for accurately mapping the small, ephemeral wetlands common in semi-arid and arid regions (Gxokwe et al., 2020). To overcome these limitations, spectral unmixing techniques have been employed to evaluate surface cover via remote sensing in arid environments (Asner and Heidebrecht, 2002; Cunnick et al., 2023) and to assess wetland evolution over time (Chang et al., 2021). Using multispectral satellite data, unmixing techniques can quantify the contributions of different surface covers to the overall reflectance spectrum of a pixel (Rasti et al., 2024). A previous study has specifically applied unmixing techniques to wetlands in dryland areas using Sentinel-2 data (Ozer and Leloglu, 2022). While this study provided a valuable method for recent wetland assessments, it was constrained by limited time frames. To address this challenge, we developed an unmixing technique based on the Landsat TM dataset, spanning from 1986 to 2021. This approach was validated using a random forest classifier trained on Sentinel-2 data.

Our research questions are the following: (1) How have water, vegetation, and soil surface covers evolved in WiD’s central and riparian zones from 1986 to 2021 in HNP? (2) What are the relationships between water, vegetation, and soil fractions in relation to hydroperiod and distance from the center of the WiDs? (3) Do temporary WiDs exhibit different surface dynamics compared to permanent WiDs?

## 2. Materials and methods

### 2.1. Study site

This study focuses on Hwange National Park (HNP), Zimbabwe. This park is characterized by a deciduous endorheic savanna and covers approximately 15,000 km^2^. The climate alternates between the dry season, from April to October, and the wet season, from November to March. During the wet season, cumulative monthly mean precipitation can reach up to 175 mm in January and drops to nearly zero between July and August (Roy et al., 2025). Temperatures have been rising since at least 1986, and drought intensity has increased across the park since 1920, although mean precipitation shows no clear trend (Chamaillé-Jammes et al., 2007a; Roy et al., 2025).

The savanna encountered in HNP, and particularly in WiDs’ vicinity, is composed of open grassland, bare soil, water, and low-density woodlands with a relatively homogeneous tree composition, depending on the geological substrate. *Colophospermum mopane* is closely associated with basaltic soil, and *Baikiaea plurijuga* is associated with Kalahari Sands (Arraut et al., 2018; Chamaillé-Jammes et al., 2009).

In HNP, water resources are available through non-perennial rivers, pans, pools, springs, and seeps (Chamaillé-Jammes et al., 2007b; Fynn et al., 2015). Their exact number is difficult to quantify due to interannual variability in precipitation and limited road access, particularly in the central part of the park (Dzinotizei et al., 2018). However, previous visual censuses identified at least 267 wetlands distributed across HNP (Figure 1). Of these, 205 are natural, most of which are temporary wetlands that may dry up by the end of the dry season (Roy et al., 2025). The remaining 62 wetlands are artificially maintained during the dry season for wildlife drinking water, using groundwater pumping. We refer to these 267 wetlands as validation points in the following (Figure 1).

**Figure 1:**
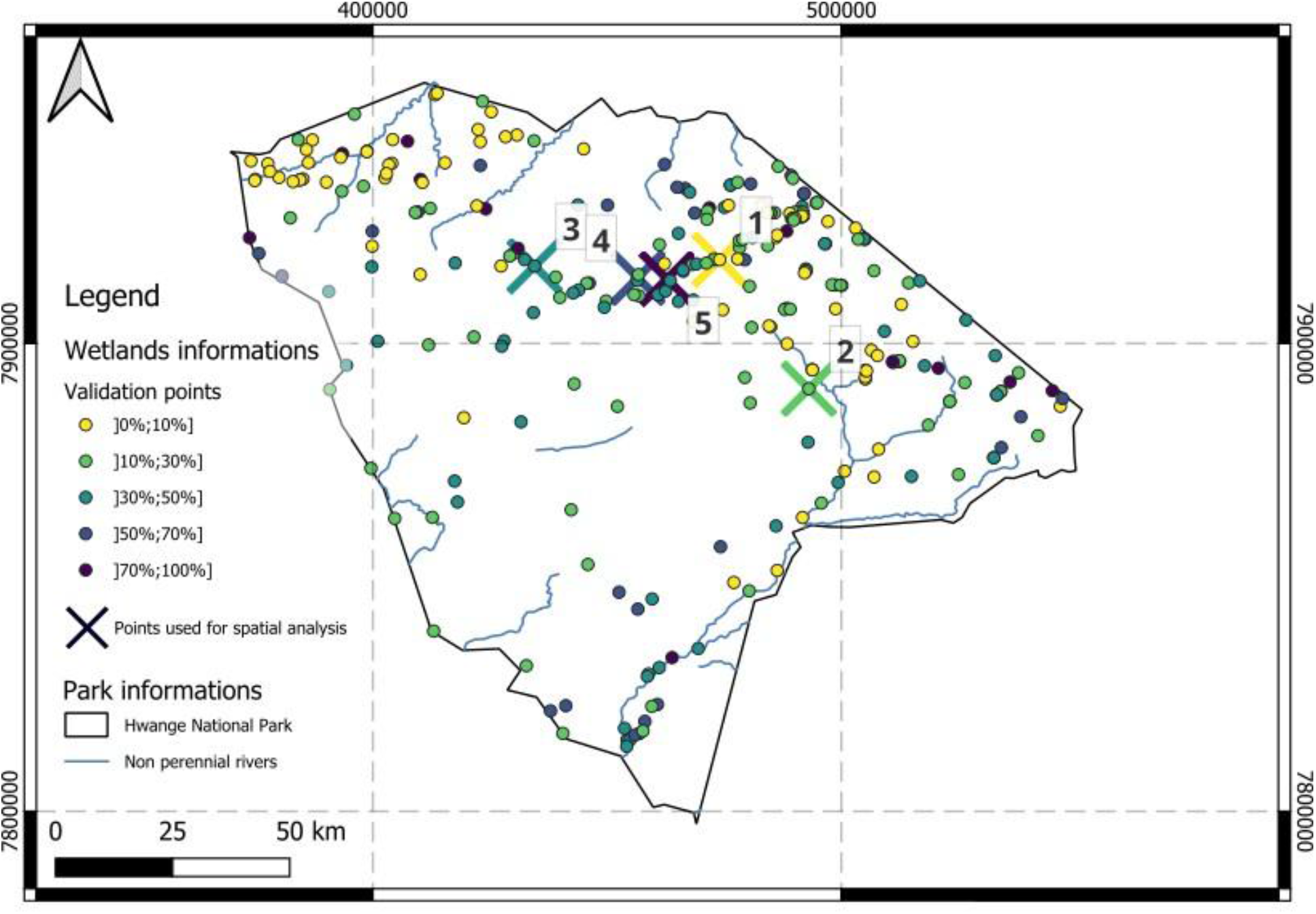
Localization of the studied wetlands in Hwange National Park. Validation points are presented witha color gradient that corresponds to water presence frequency (See section 2.2.1). Crossed points are used for spatial visual analysis (see Section 2.2.4). Their respective names are 1) Pan Urge, 2) Togo, 3) Roan pan, 4) Shapi, 5) White Hill. The coordinate reference system is ESPG: 32735 - UTM zone 35S.

### 2.2. Remote sensing data extraction and unmixing

#### 2.2.1. WiD’s ephemerality assessment and attribution

To assess the ephemerality of WiDs, water presence frequencies were calculated with a Thresholding Single Water Index method using Landsat 5, 7, and 8, with data ranging from 1986 to 2022 (Roy et al., 2025). Water detection maps were extracted using scripts written in Google Earth Engine (Gorelick et al., 2017) and Landsat collections as inputs. These methods are described below.

Water detection is performed using the Modified Normalized Difference Water Index (MNDWI). It is calculated as the normalized difference between the green band and the shortwave infrared (SWIR) band of Landsat (Xu, 2006). The green band corresponds to reflectance in between the 0.533 and 0.590 µm wavelengths, while the SWIR band corresponds to reflectance in the wavelengths between 1.566 and 1.651 µm.

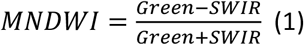

MNDWI varies from -1 to 1. A higher contribution of the green band, which is linked to the presence of water, raises the index. The MNDWI threshold is set at -0.29, which was previously defined as optimal for the HNP region (Dzinotizei et al., 2018). All index values equal to or above this threshold indicate the presence of water within the pixel.

For each image extracted from Landsat, every pixel is given a value of 1 if water is detected, and 0 in the absence of water. The frequency of water availability (i.e., the frequency of water detection) at a given pixel is calculated for the whole period (2).

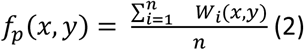

*f*_*p*_(*x, y*): probability of water presence for the pixel at location (*x, y*) for the considered period *p* (here from 1986-2022).

*W*_*i*_(*x, y*) : presence of water at time *i* at location (*x, y*). Equal to 1 if water is present and 0 if water is absent.

*n*: number of times the pixel is observed for the period 1986-2022.

For each of the 267 validation points, the water frequency of the overlapping pixel was extracted and assigned to the corresponding wetland. It is used to evaluate the water availability of the 267 known wetlands by providing a frequency of water presence in HNP. After extracting this data (Roy et al., 2025b), we classified each WiD based on its overall frequency of water presence, thereby creating a categorical classification of hydroperiod. The following classes are settled: [0%,10%], [10%,30%], [30%,50%], [50%,70%], [70%,100%]. The number of WiDs in each frequency class is shown in Figure 2.

**Figure 2:**
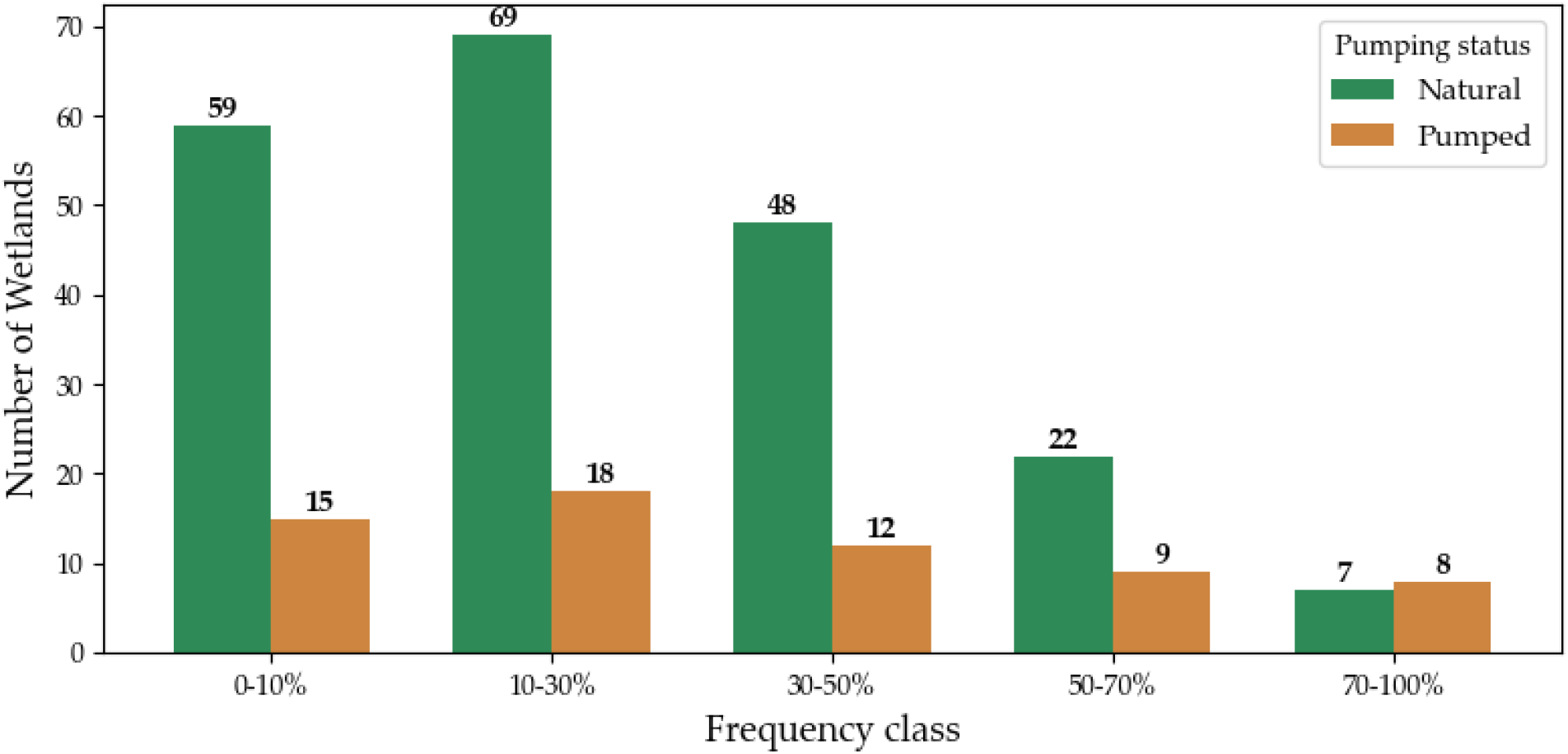
Proportion of WiDs in each frequency class. Natural wetlands are represented in green, pumped wetlands are in brown. The number in each class is mentioned above the bar.

We hypothesize that water frequency is the primary driver of riparian variations. Although we have illustrated the repartition between pumped and natural WiDs in Figure 2, we do not distinguish them in the following. Indeed, a previous study showed that the pumping status does not influence the size of the piosphere gradient, leading to similar degradation in pumped and non-pumped WiDs with similar hydroperiods (Roy, 2025).

#### 2.2.2. Data extraction and processing

The processing and extraction of remote sensing satellite images have been carried out using the Google Earth Engine (GEE) cloud-based platform (Gorelick et al., 2017). As shown in Table 2, the Landsat time series we used covers the period from 1985 to 2022. We extracted data from Landsat collections (LANDSAT/LT05/C02/T1_L2, LANDSAT/LE07/C02/T1_L2 LANDSAT/LC08/C02/T1_L2) and Sentinel-2 collection (COPERNICUS/S2_SR_HARMONIZED) for validation purposes.

**Table 1:** Collection of images extracted in Google Earth Engine for the unmixing and random forest methods.

| Satellite name | Collection of images in GEE | Dates used |
| --- | --- | --- |
| Landsat 5 | LANDSAT/LT05/C02/T1_L2 | 1985.01.01 - 2011.12.31 |
| Landsat 7 | LANDSAT/LE07/C02/T1_L2 | 2012.01.01 - 2013.02.28 |
| Landsat 8 | LANDSAT/LC08/C02/T1_L2 | 2013.01.01 - 2022.12.31 |
| Sentinel-2 | COPERNICUS/S2_SR_HARMONIZED | 2018.10.01 - 2020.03.31 |

Images extracted from GEE were filtered at the pixel scale using quality assessment flags to mask cloudy or shadowed pixels.

Mean monthly 2m air temperature data were extracted from ERA5 (ECMWF/ERA5/MONTHLY) (Muñoz-Sabater et al., 2021) for the park area, covering the period from January 1985 to June 2020, at a spatial resolution of 30 km. The Normalized Difference Vegetation Index (NDVI) is extracted for each Landsat pixel.

#### 2.2.3. Intra-pixel spectral unmixing of the local environments surrounding water sources, at the Landsat pixel scale

To properly evaluate the surface cover evolution around WiDs and obtain sufficient data in a timely manner, the Landsat project appears to be the most suitable satellite project. However, the main drawback of Landsat is its spatial resolution of 30m, whereas many WiDs, especially in HNP, are smaller than this pixel size (Roy et al., 2025). On the other hand, Sentinel-2 multispectral satellite provides images with a resolution of 10m, which would greatly enhance the spatial resolution compared to Landsat. But this gain comes with a short data availability, from 2018, weakening the potential temporal trend, which may be observable with Landsat’s operation timespan. To overcome this trade-off, we perform a linear multispectral unmixing (LMU), detailed below.

Each pixel in a satellite image captures a mixture of different materials, and spectral unmixing aims to decompose this mixture into pure spectral signatures, or endmembers, and their respective abundances within the pixel. The complexity of the unmixing process increases with the number of endmembers and the required precision, as each additional endmember introduces a new variable to the linear mixing model, thereby making the system of equations more complex to solve accurately. In this study, we aim to differentiate between water, bare soil, and vegetation within the same Landsat pixel. These categories present contrasting optical properties, and several unmixing studies have demonstrated the feasibility of identifying signals specific to each element (Asner and Lobell, 2000; Guerschman et al., 2015). Using the linear unmixing approach is an approximation. Indeed, there are linear and multilinear (e.g., bilinear) models for spectral unmixing. The linear approach assumes that endmembers are mixed linearly, which is valid when each light ray interacts with only one material before reaching the sensor. This assumption is commonly used in macroscopic questions, such as those related to Earth observation, but it is inherently an approximation. Bilinear mixing assumes that light interacts with two materials before reaching the sensor, leading to double scattering (Rasti et al., 2024). In Earth observation, non-linear behaviors can be induced by various factors, such as noise, atmospheric effects, temporal effects, illumination variations, and the intrinsic spectral variability of materials. For instance, changes in material moisture content or seasonal variations in leaf spectral signatures can significantly affect spectral data. The difference between supervised and semi-supervised unmixing lies in how the endmember information is provided. In supervised unmixing, the model is trained using known spectral signatures, whereas in semi-supervised unmixing, the model is provided with a library of endmembers to choose from.

Because of the low density of woodlands in the park, and the rather open savannah structure (Arraut et al., 2018), the risk of bilinear mixture and double scattering is low. Furthermore, linear models are often sufficient for Earth observation questions with atmospheric correction (Rasti et al., 2024). Given these elements, we opted for supervised unmixing within the linear mixing model framework.

Three endmembers were defined from Landsat images to represent the spectral signatures of water, vegetation, and bare soil (Figure 3). Even within a year, vegetation presence and its spectral signature vary significantly due to high intra-annual climatic variability (Almalki et al., 2022; Chamaille-Jammes et al., 2006; Nicholson and Farrar, 1994; Schmidt and Karnieli, 2000). To correctly describe the vegetation patterns, our work focused solely on the wet season. This allows for the correct evaluation of vegetation surface evolution over long periods. Mean images were calculated for each wet season, spanning from November 1st to March 31st of the following year. Here, each wet season is named after the year in which it ends.

**Figure 3:**
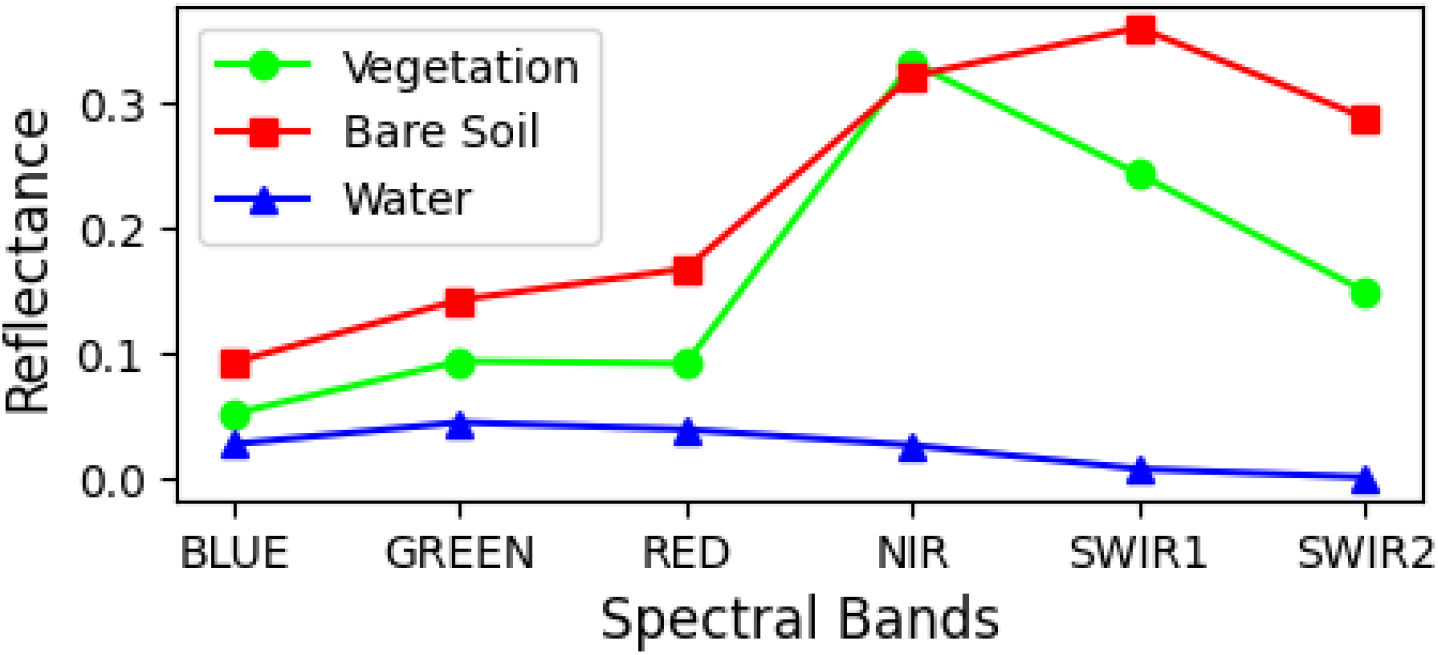
Spectral signature of each endmember used for the unmixing classification.

Each wet season mean image was unmixed using the .unmix() function available in Google Earth Engine (GEE). The spectral signature of each endmember was used to decompose the pixel reflectance into the proportion contributed by each land cover type, thus estimating the percentage of surface covered by water, vegetation, or bare soil for each wet season.

To contrast the reliability of the LMU carried out with Landsat, Sentinel imagery was used. Since it provides better pixel resolution (10×10 m instead of 30×30 m), a surface classification was performed to contrast with the unmixed images. Mean wet season Sentinel-2 (S2) images of the years 2019 and 2020 were classified into: Free water bodies, Wetlands, Bare soils, Herbaceous crops, and Shrubs/Trees. Training polygons were chosen based on the ecological specificities of HNP. Free water bodies were identified and used for training. Wetlands were also used as training polygons to incorporate the specific features of such an ecosystem, including water, vegetation, and bare soil, with specific variability. For vegetation, we used crop polygons as training. Although no crops are found in HNP, this choice was motivated by the challenge of locating training polygons that exclusively represent herbaceous surface cover, devoid of bare soil or trees. Selecting crops was the only viable option in the region to ensure proper extraction of vegetation features by the random forest. For this purpose, some training polygons were selected visually from Landsat, map, and Google Earth images for each random forest (RF) classification output, and an RF supervised machine learning process was implemented using 25 trees. The training sample (80% of data) has an accuracy of 0.999, and the validation sample (20% of data) has an accuracy of 0.995. The validation confusion matrix is available in Supplementary Materials (Table S1).

#### 2.2.4. Analysis of the spatio-temporal variation of NDVI in the vicinity of waterholes

To analyze NDVI variations in the surroundings of the wetlands at different distances, summary statistics of NDVI were retrieved from concentric circles (more precisely, donut-shaped buffers) of varying sizes around each wetland. These buffers, centered on locations marked in the wetland validation dataset, had radii ranging from 50 m to 400 m, in 50 m increments. For each wet season and each wetland, the mean value (as well as the standard deviation and the number of pixels within the buffer) is retrieved at each distance for the vegetation, soil, and water fractions as well as for the NDVI. The total surface area of all pixels located at the given radius and in all directions is taken into account. The images were processed using QGIS.

To assess the accuracy of Landsat intra-pixel unmixing results, the fractional compositions of the three land cover categories were compared with land cover classifications derived from Sentinel-2 imagery for the same season in the years with available data (2019–2020). The same method, based on the use of identical buffers, was applied to extract the relative composition of land cover within each buffer using random forest classifications of Sentinel-2 images as inputs. Each land cover category’s proportion from S2 classifications was calculated by counting the number of pixels assigned to that category within the same buffer zone.

Time series of land cover fractions and NDVI are constructed from 1986 to 2022. The overall process is presented in Figure 4.

**Figure 4:**
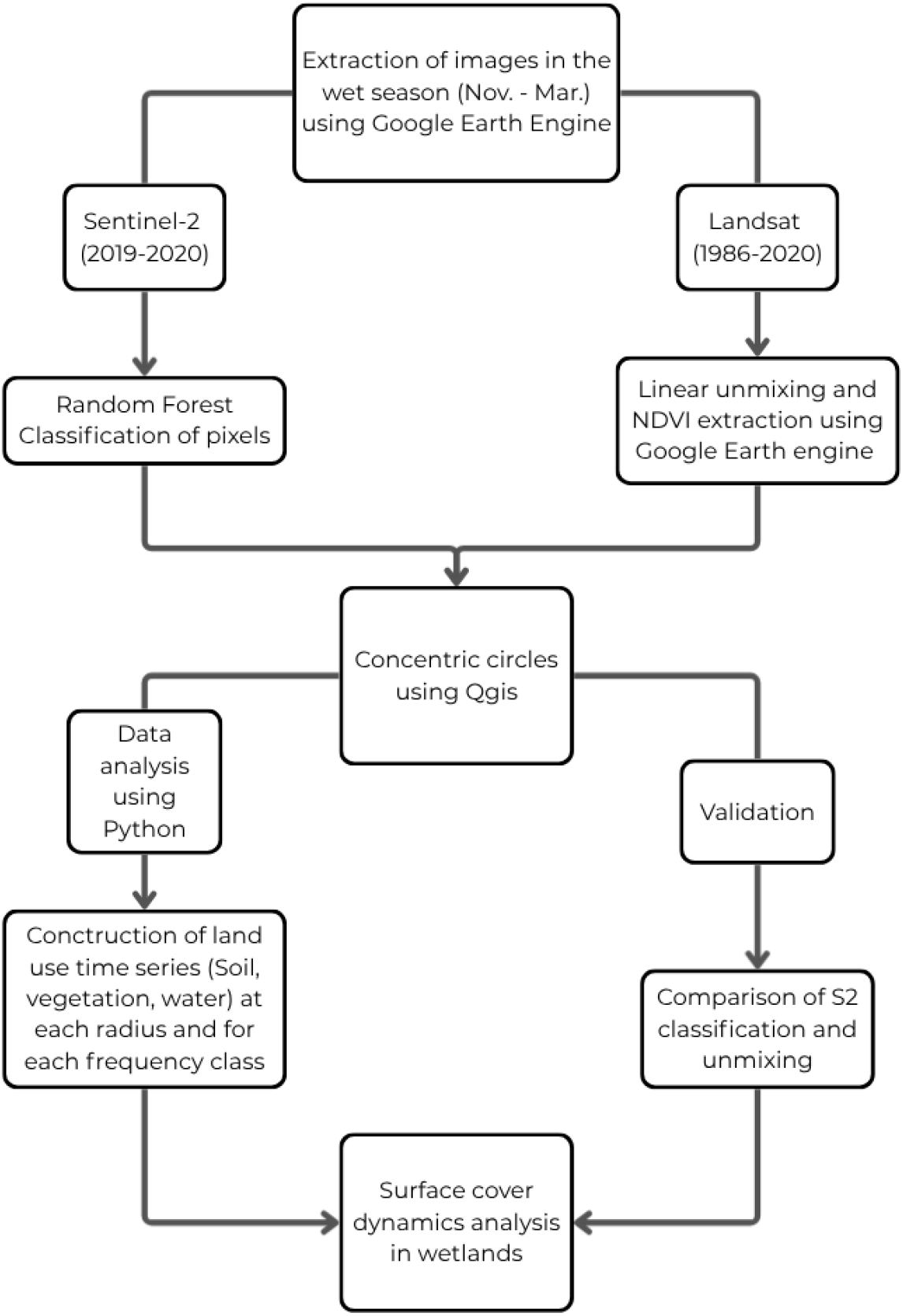
Conceptual diagram of the image analysis methodology.

#### 2.2.4. Statistical analysis

The land cover fractions derived from Landsat and Sentinel-2 were compared using regression lines, R-squared, and root-mean-square error (RMSE).

Time series of land cover fractions were constructed for each distance. The validation was performed on the 200m wide buffer, which is half the maximum buffer, to represent a compromise between distal and proximal wetland dynamics.

Linear regressions are performed for each distance and frequency class to assess trends in water, bare soil, and vegetation cover over time. Kendall’s tau correlation coefficients are calculated between the different land cover fractions at each distance and frequency class. Additionally, Kendall correlations are computed between each fraction and temperature, at each distance, and for each frequency class. Correlations with precipitation were also assessed, but did not exhibit significant results and are therefore not presented.

To ensure that the observed trends reflect actual changes rather than temporal dependencies, the autocorrelation of the detrended water fraction was assessed at lags 0 to 4. The results, presented in the Supplementary materials : Table S2, show no significant autocorrelation, except for a positive autocorrelation in the water fraction of wetlands within the [0–10%] frequency class.

## 3. Results

### 3.1. Unmixing validation

Figures 5 and 6 show the comparison between the fraction estimates from the LMU and two distinct validation metrics.

**Figure 5:**
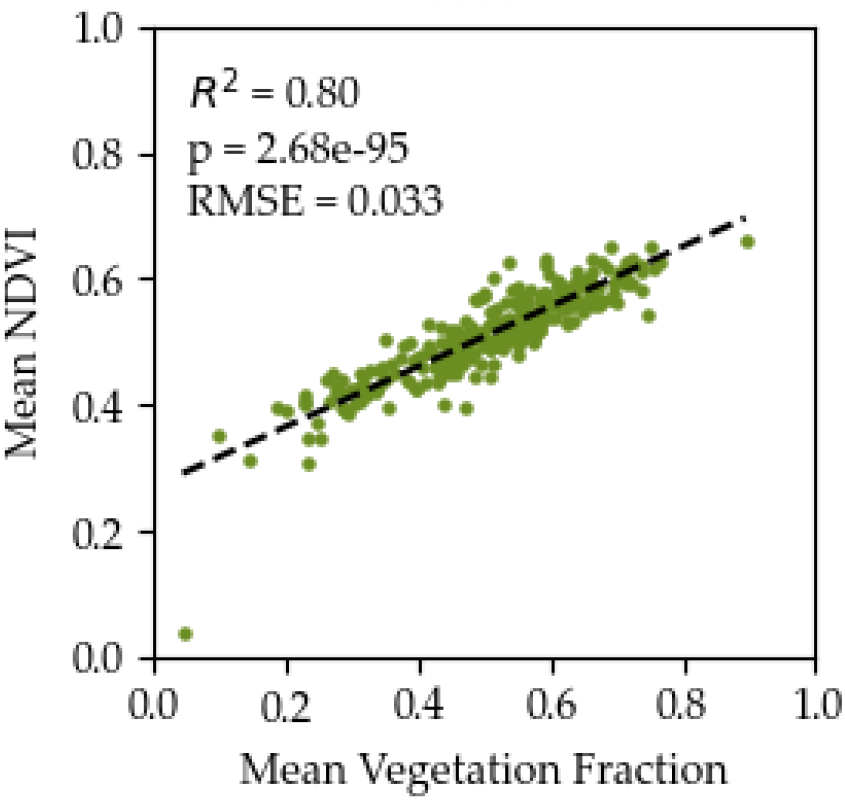
Validation of the vegetation fraction extracted from the LMU with the NDVI in year 2021. R^2^, p-value, and RMSE are mentioned in the plot.

**Figure 6:**
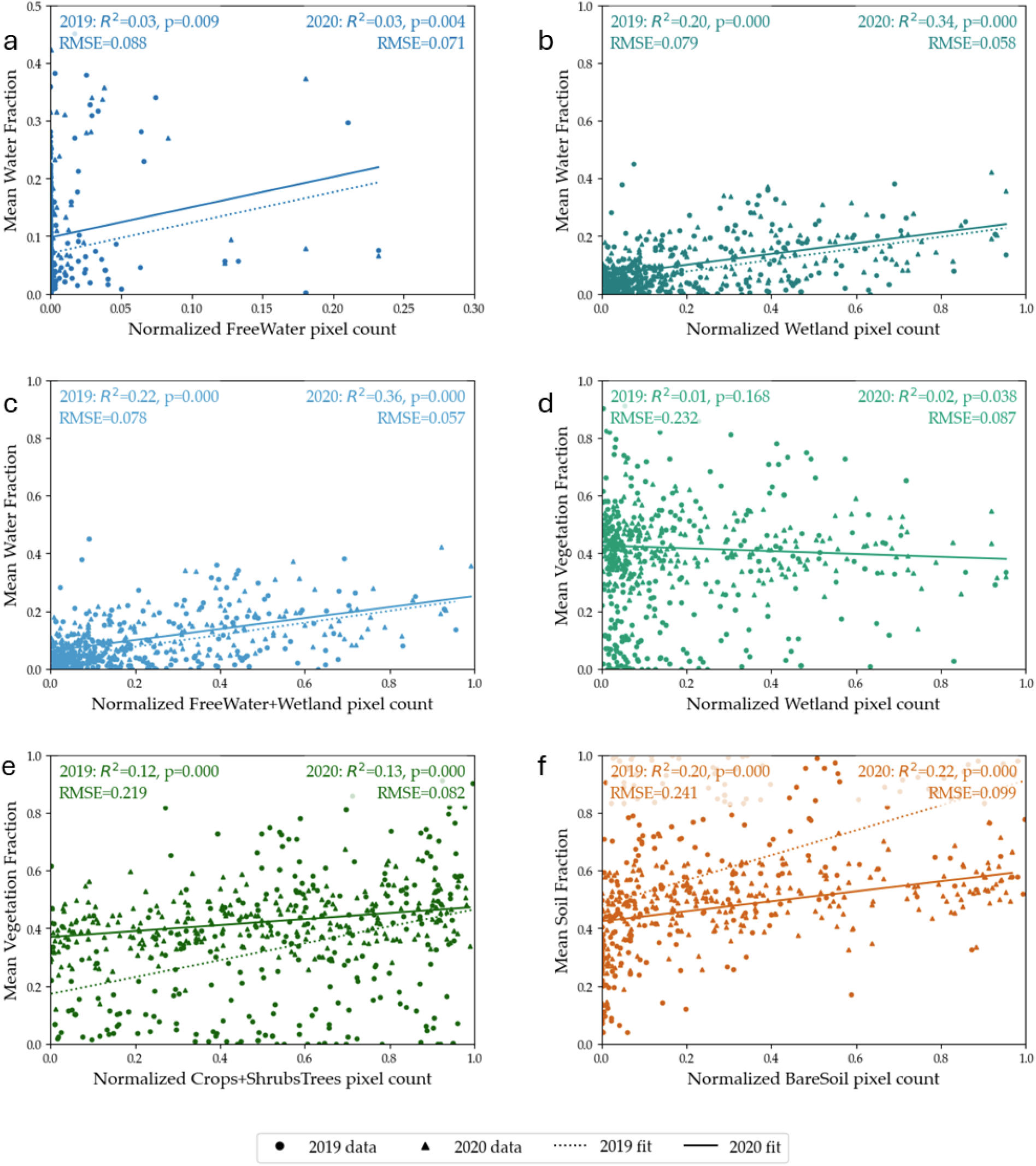
Validation of the Linear Multispectral Unmixing with a random forest Sentinel 2 model. Round points and triangles represents respectively 2019 and 2020 data. Dotted and continuous lines represent respectively the 2019 and 2020 significant fit. R^2^, p-value, and RMSE for each year are mentioned on each graph. The axes are adapted to the range of data. a. Correlation between the water fraction and the normalized free water pixel count. b. Correlation between the mean water fraction and the normalized wetland pixel count. c. Correlation between the mean water fraction and the normalized sum of free water and wetland pixel count. d. Correlation between the vegetation fraction and the normalized wetland pixel count. e. Correlation between the vegetation fraction and the normalized sum of the crops and shrub tree pixel count. f. Correlation between the normalized soil fraction and the bare soil pixel count.

The linear regressions between NDVI and vegetation fractions each year from 1986 to 2021 (which includes the beginning of 2022) are highly significant (Figure 5, Supplementary Materials: Figure S1). For instance, in 2021, the R^2^ is 0.80 and the RMSE is 0.033.

A comparison of S2 RF classification pixel counts and LMU results revealed a significant correlation between almost every fraction and its corresponding RF classification outcomes for both 2019 and 2020 (Figure 6). One notable exception was observed in the relationship between wetland pixel counts and vegetation fraction in 2019, where the correlation is not significant (Figure 6d). Every other regression was significant (p<0.05) to highly significant (p<0.005). Additionally, the combination of normalized crops and shrub trees demonstrated a highly significant correlation with the vegetation fraction (Figure 6e). For the water fraction, the R^2^ value generally improved when pixel counts were associated with classified pixels of both free water and wetlands (Figures 6a and 6c). The RMSE was overall higher in 2019 compared to 2020, with a maximum value of 0.241 for the Mean Soil Fraction against Bare soil pixel count (Figure 6f). The RMSE was inferior to 0.1 for the Water fraction against the Free water and the wetland pixel count (Figure 6c).

### 3.2. Surface cover dynamics

The evolution of the three fractions in each spatial buffer is shown in Figure 7. The vegetation and soil fractions exhibit almost symmetrical evolution through the time series. A local maximum in vegetation fraction is simultaneous with a local minimum in soil fraction and vice-versa. Between 2003 and 2005, these two fractions exhibited distinct opposite patterns at each distance. The vegetation fraction occupies between 0 and 60% of the surface area, whereas the soil fraction occupies between approximately 20 and 90% of the surface area. The remainder is occupied by water (between 5 and 30%). The water fraction is significantly decreasing from 150m to 400m. The linear regression is significant from 150m to 250m (*p<0*.*05)* and highly significant from 300m to 400m (*p<0*.*005)*. The slope of the regression increases with distance, ranging from -0.0014 at 150m to - 0.0017 at 300m to 400m.

**Figure 7:**
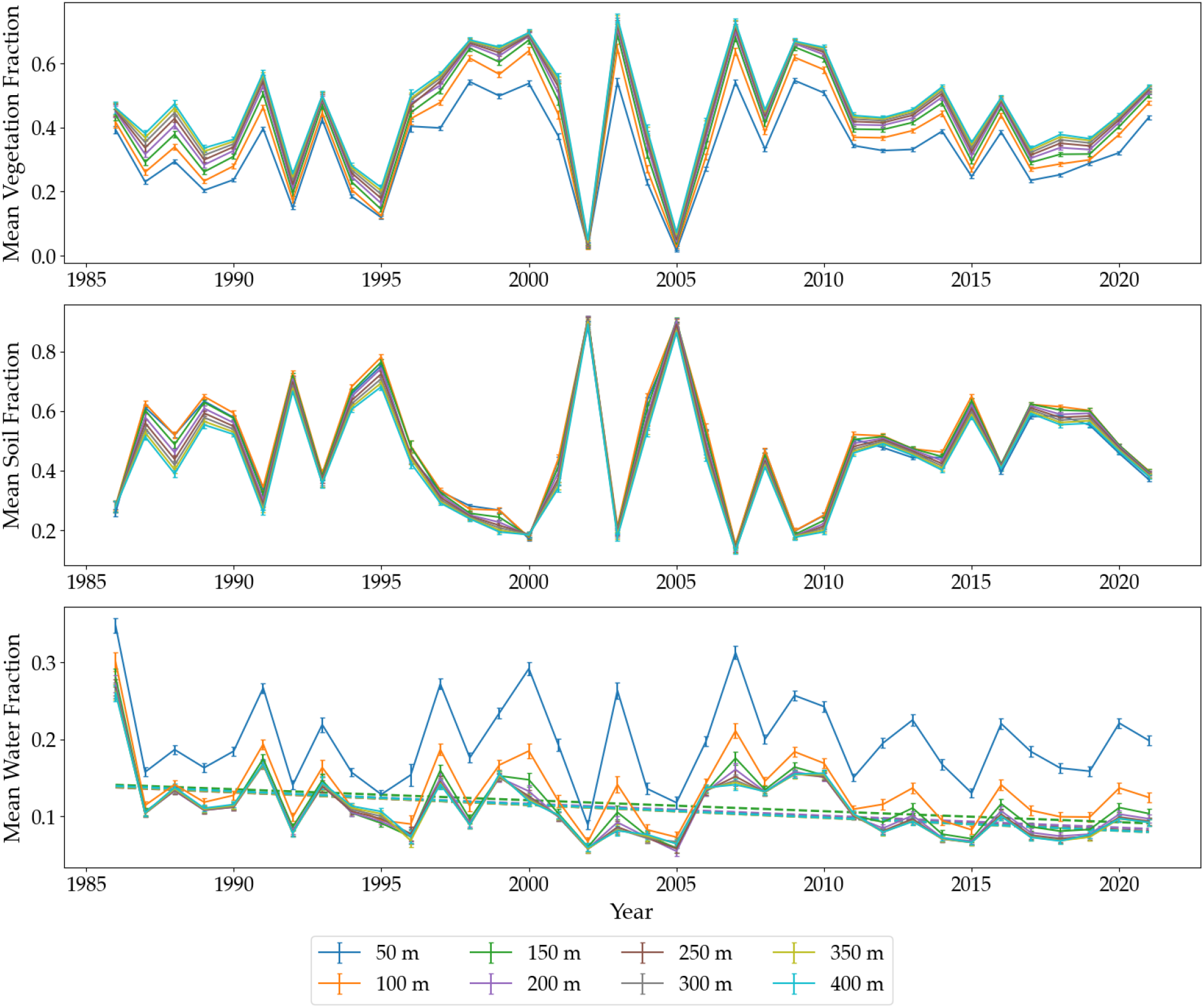
Evolution of vegetation, water and soil fractions from 1986 to 2021 at each distance from the WiD’s center. Significant linear regressions are shown as dashed lines.

The vegetation fraction is higher for classes with lower water frequency, whereas the soil fraction exhibits the opposite pattern, with higher fractions corresponding to higher water frequency (Figure 8). Regarding their dynamics, the vegetation and bare soil fractions exhibit antagonistic behavior, even within a single water frequency class. The linear regression with time for these two fractions is not significant. The water fraction significantly decreases from classes 0-10% to 30-50%. The results are highly significant for the 0-10% class (p < 0.005) and significant for the 10-30% and 30-50% classes (p < 0.05). The slope decreases as frequency classes increase, ranging from -0.0019 for the 0-10% class to -0.0013 for the 30-50% class. All classes show the presence of water in every wet season, whereas the vegetation fraction reaches 0 in 2003 and 2005 in every class.

**Figure 8:**
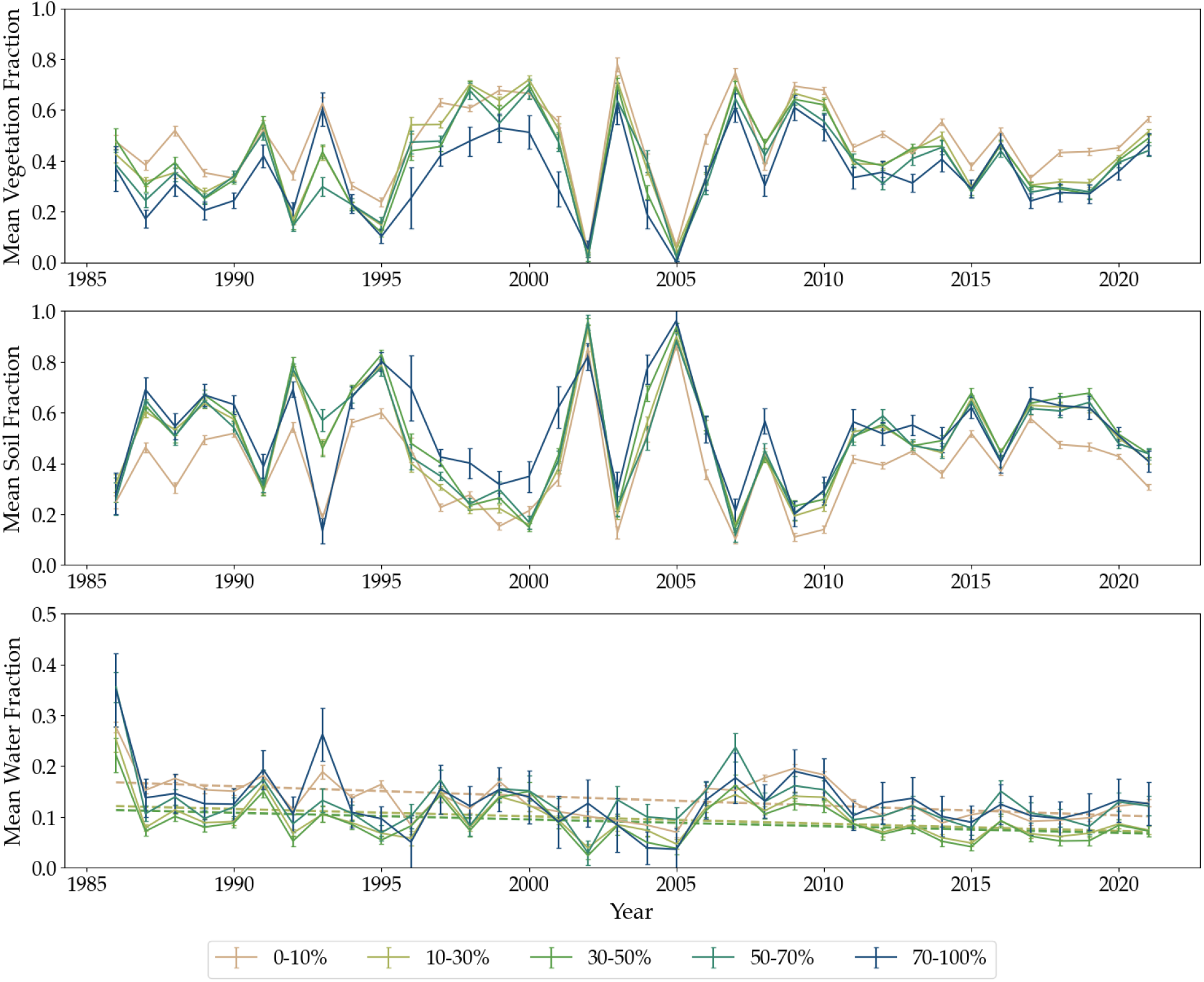
Evolution of each fraction from 1986 to 2021, depending on the frequency of water availability. This calculation is based on a 200m buffer. Significant linear regressions are shown as dashed lines.

### 3.3. Fractions inter-dependance and link with climate

The Kendall correlation between vegetation and water fraction is always positive along the buffer radius, decreasing from 50 m to 200 m and reaching a minimum of 0.333 at 300-400 m (Figure 9a). Correlation between soil and water exhibits an opposite pattern, with an increasing negative correlation, starting from -0.730 at 50m to -0.440 at 400m. The correlation between vegetation and soil remains stable at every distance, with a value of approximately -0.9. Along the frequency class gradient, vegetation and water are still positively correlated (Figure 9b). Vegetation and soil, on the one hand, and soil and water, on the other hand, remain negatively correlated. The correlation between vegetation and water increases from 0.197 for the 0-10% class to 0.521 for the 30-50% class. The correlation between soil and water shows an opposite pattern, decreasing from -0.337 for the 0-10% class to -0.597 for the 30-50% class. The correlation between vegetation and soil remains stable across all classes, at approximately -0.9.

**Figure 9:**
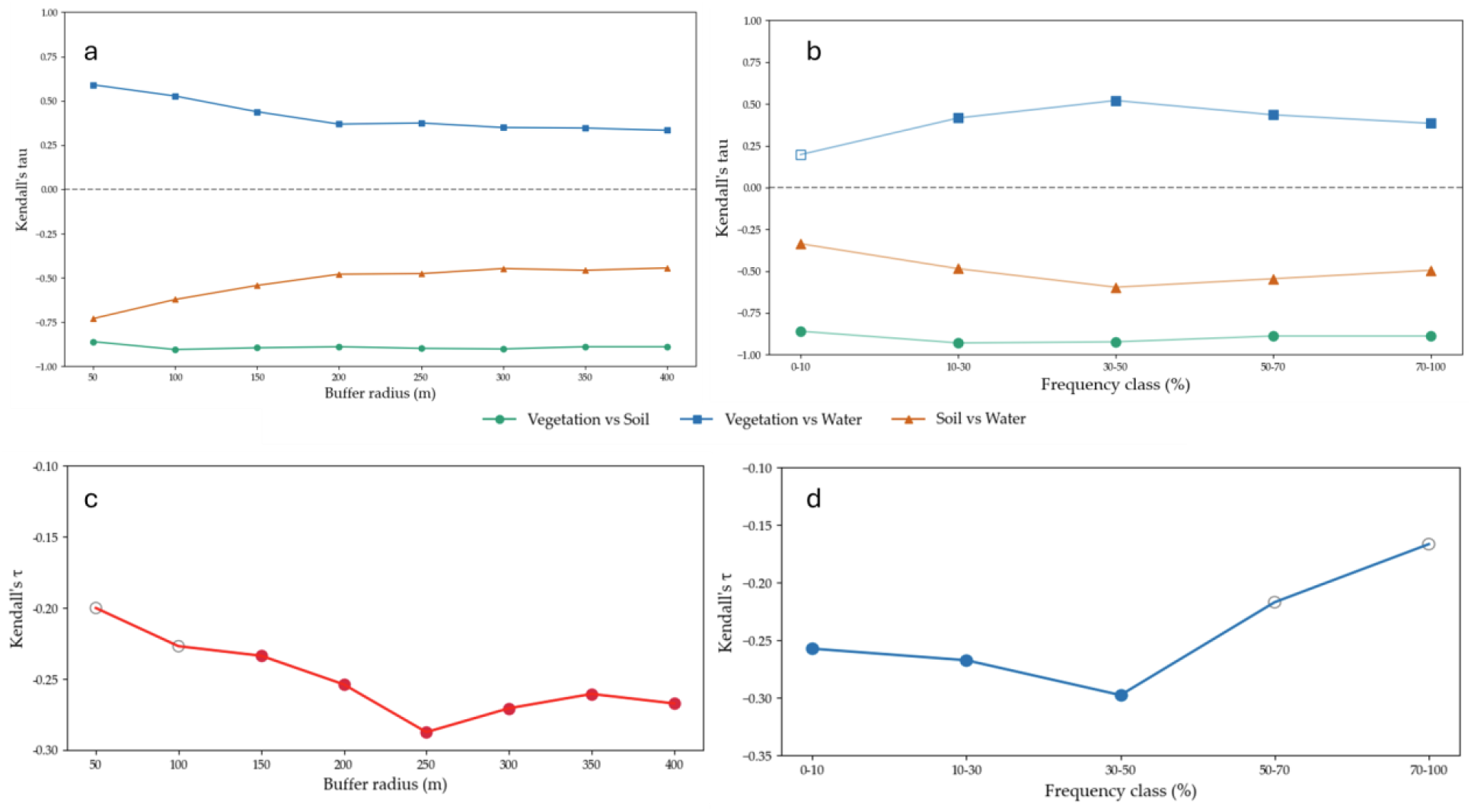
Kendall correlation between fractions along the distance gradient (a) and the frequency of water availability gradient (b). Kendall correlation between water fraction and temperature as a function of the distance gradient (c) and the water frequency class (d). Full dots represent significant correlations. Emptied dots represent non-significant correlations.

The Kendall correlation between temperature and water fraction was significantly negative from a 150m to a 400m buffer radius (Figure 9c). It remains relatively stable, with a mean of -0.262, a maximum of -0.227 at 150m, and a minimum of -0.287 at 250m. Kendall correlation between temperature and water fraction is significantly negative from the 0-10% to 30-50% class of water frequency (Figure 9d). It exhibits a decreasing pattern, with a maximum of -0.257 for the 0-10% class, down to -0.297 for the 30-50% class.

Figure 10 presents visual examples of WiDs from each water frequency class, illustrating their dynamics across different years. The impact of inter-annual climate variability is clearly visible. The dry year of 2005 stands out, with nearly total vegetation disappearance. During this year, only the more frequent WiDs showed a few remaining pixels of water, whereas the water fraction at Pan Urge was not visible at all. In contrast, wet years such as 2000 and 2020 show much more extensive water surfaces. While all sites are affected by this climate variability, they respond differently according to their frequency class. Water is visible only in wet years for Pan Urge (0%-10% class), where its spatial repartition was also more heterogeneous and seemed less spatially concentrated, especially in 2000. Conversely, water is always available in White Hill (70%-100% class), even in the dry year of 2005. A key pattern is the presence of bare soil in the immediate vicinity of the water. For more frequent WiDs, such as Roan Pan (30%-50% class) and White Hill, the presence of bare soil in the immediate vicinity is particularly visible throughout the time span of the figure. The Roan Pan example from the wet year of 2000 suggests that this bare soil proximal surface cover could correspond to a potential maximum inundation surface. In contrast, this clear bare soil fraction in the water’s immediate vicinity is not observable for the ephemeral Pan Urge.

**Figure 10:**
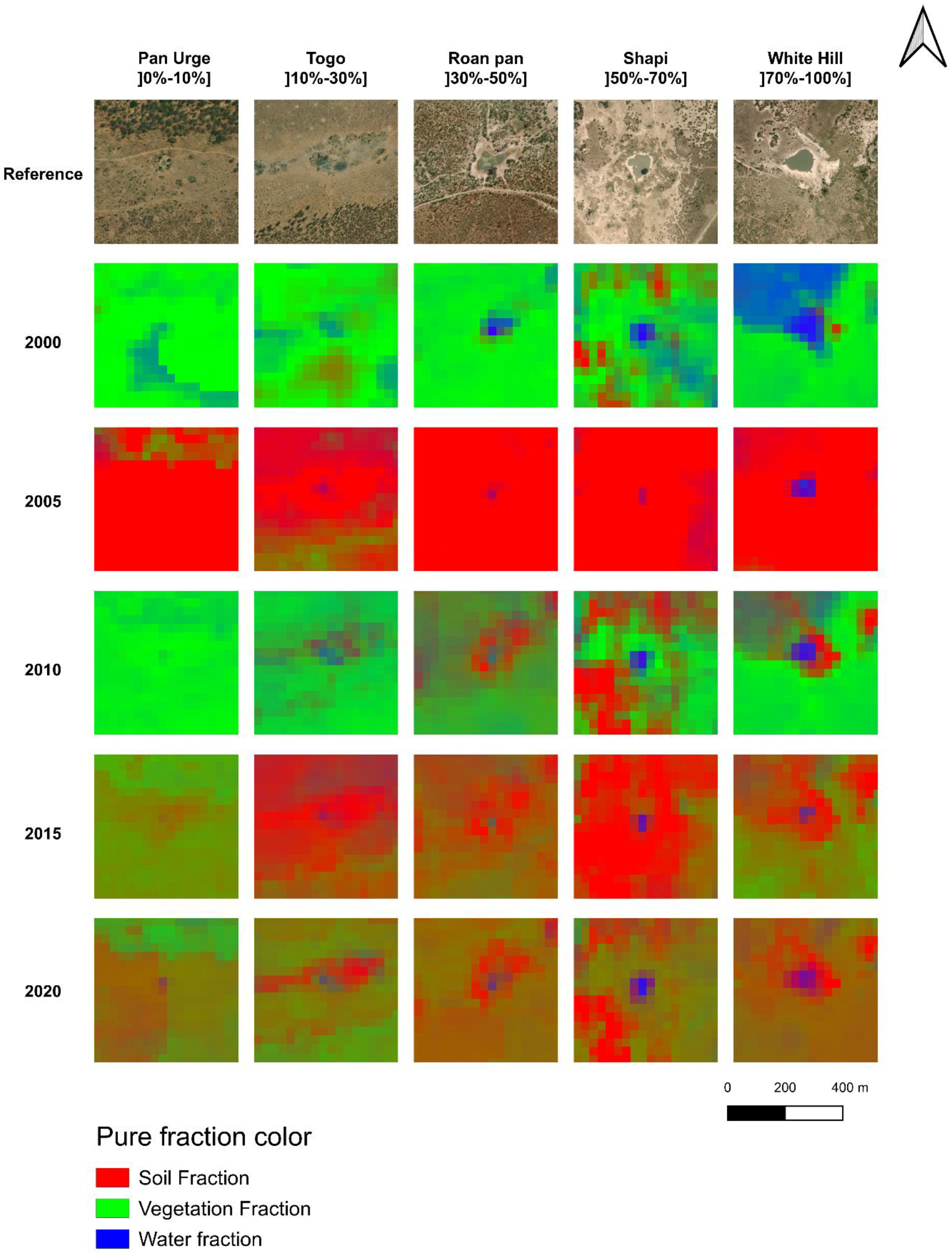
Mean surface cover evolution from the 2000 to the 2020 wet seasons in 5 WiDs from each water frequency class.

## 4. Discussion

### 4.1. Validation of the Linear Multispectral Unmixing method against Random Forest classification

The LMU demonstrated a strong capacity to detect each fraction when compared to a highly accurate RF classification. Furthermore, the vegetation fraction was significantly correlated with NDVI, reinforcing confidence in the precise estimation of vegetation cover and biomass.

The water fraction was significantly correlated with the free water pixel count in both validation years; however, the R^2^ was low (0.03 in both years). A stronger correlation was observed between the water fraction and the wetland pixel count, with higher R^2^ values (0.20 and 0.33 in 2019 and 2020 respectively). The best agreement was observed when the water fraction was compared with the addition of water and wetland pixels, yielding the highest R^2^ (0.22 and 0.36). In contrast, the vegetation fraction was significantly correlated only with the wetland pixel count in 2020, and even then, with a low R^2^ (0.02). This may be explained by the nature of the wetland training polygons used for the RF classification, which included both permanent water and adjacent riparian zones that can be seasonally flooded and covered by vegetation or bare soil during other periods. Moreover, substantial interannual climatic and hydrological variability (Roy et al., 2025) likely affects the optical signature of wetlands across years. Overall, our results suggest that, based on the selected RF training polygons, the LMU-derived water fraction effectively captures the optical response of wetland areas, and to a lesser extent, free-standing water.

Vegetation fraction was significantly correlated with the addition of crop and shrub/tree pixel counts in both 2019 and 2020, demonstrating that it effectively captures the proportion of vegetation within a pixel. A similar pattern was observed between the soil fraction and bare soil pixel counts. Although the correlations were highly significant (p<0.005), they raise questions about the LMU approach’s ability to accurately represent the actual surface composition of both fractions. Nevertheless, the vegetation fraction was also strongly correlated with NDVI across all years, with a minimum R^2^ of 0.26; 1986 was a year with particularly low vegetation cover in the park (see Figure S1). This supports the conclusion that the vegetation fraction is an effective proxy for vegetation biomass. Indeed, the reference polygons used, particularly for crops, were chosen for their optical homogeneity, which is harder to achieve in natural dryland systems, where grasslands are often heterogeneous mosaics of vegetation patches, soil, and scattered trees. This explains both the vegetation fraction’s ability to capture vegetation biomass and the high error rates in the S2-based random forest (RF) classification.

Similar limitations affect the correlation between soil fraction and bare soil, as bare soil polygons may contain some vegetation, contributing optical signals that lead the RF to classify pixels as bare soil even if vegetation can be found inside. The LMU, by design, does not have this problem and quantifies each fraction inside the pixel. These challenges stem from the inherent complexity of the natural landscapes studied.

Overall, the LMU classification accurately captures the spatial and temporal dynamics of wetlands, vegetation, and bare soil. While some errors likely result from limitations in classification design and spatial resolution, the strong correlation with NDVI provides confidence in LMU’s ability to estimate vegetation biomass. However, these potential errors should be taken into account when using the model for precise surface quantification.

### 4.2. Hydroperiod and its buffering role against climate change

Our results showed that the water surface significantly decreased from 1986 to 2020, at distances ranging from 150m to 400m. We observed the same decreasing pattern for temporary wetlands with water frequencies ranging from 0% to 50%. Therefore, we demonstrate that the hydroperiod mitigates wetland vulnerability, as previously known for temperate wetlands (Boix et al., 2020) but not for dryland wetlands, to the best of our knowledge.

The water fraction reached its highest value in 1986. We verified the acquisition conditions for this year: images were collected in November–December, months typically associated with lower water availability during the wet season (Roy et al., 2025). Although these months did not reach peak wet-season levels, water availability was 18–30% above the November–December average, consistent with interannual variability rather than anomalous conditions. The 1986 data point, therefore, reflects actual hydrological conditions and was retained in the analysis.

The decrease in water surfaces was noticeable at distances of 150-400m from the center of the WiDs. Variability in water levels was greater within buffers smaller than 150m, as water contributed more significantly to the overall surface. This increased variability may mask long-term signals, which become more evident with wider buffers, where fluctuations are reduced by the contribution of vegetation and bare soil. This assertion is supported by the observed increase in slope with larger buffer sizes, indicating that a decrease also occurs in smaller buffers, albeit to a lesser extent. Therefore, the surface area of wetlands in HNP has decreased from 1986 to 2020. This pattern of wetland surface reduction is also observed in Chinese semi-arid climates, underlying a potentially consistent pattern in semi-arid drylands (Wu et al., 2024; Xue et al., 2024). This finding is also consistent with future projections for Southern African inland wetlands, which are expected to decrease in size under both RCP 2.6 and RCP 8.5 (Xi et al., 2021).

A former study showed that the longer the hydroperiod, the larger and more intense the piosphere (Roy, 2025). Therefore, the link between the hydroperiod and the ecological state of wetlands is complex. A longer hydroperiod leads to less vulnerability to climate change but a larger, more intense piosphere, whereas a shorter hydroperiod results in greater water surface loss and a smaller, less intense piosphere. Hydroperiod thus plays a dual role: enhancing hydrological resistance while simultaneously intensifying landscape degradation. Furthermore, by modifying the hydroperiod (Roy et al., 2025), pumping will move this dual role of a single wetland.

The analysis of water frequency correlation on each fraction was performed on the 200m buffer. This choice was motivated by the need to balance the contribution of each fraction within the buffer. A larger buffer would have reduced water variability, while disproportionately increasing the influence of vegetation and bare soil, potentially masking long-term tendencies. At this distance, the water fraction decreases significantly for the wetlands with a frequency of water presence below 50%. This slope increased with the frequency class, indicating that the more temporary wetlands were experiencing a more substantial decrease in their water fraction. The classes in the 0%-50% range were also significantly correlated to temperature, which is increasing in the park (Roy et al., 2025). Therefore, it confirmed that temporary wetlands are more threatened by climate change than permanent ones. This finding is consistent with previous results, which show that temporary wetlands are more threatened by temperature increases, whereas permanent wetlands tend to reduce their hydroperiod rather than their size (Boix et al., 2020; Parra et al., 2021; Sandi et al., 2020).

### 4.3. Relationship between the water, vegetation, and bare soil and its implications

Vegetation and bare soil do not exhibit any long-term tendency, unlike water, which has shrunk. However, water fraction is correlated with the other fractions. Studying how each fraction varies relative to the others is essential to understanding the process and its dynamics.

As expected in drylands, where water availability is a critical factor for vegetation growth, our analysis confirms that vegetation and water fractions are positively correlated, consistent with the literature (Chamaille-Jammes et al., 2006; Zhao et al., 2021; Zhou et al., 2021). Vegetation shows no long-term significant trend. However, its correlation with water, which is significantly decreasing in temporary wetlands, may be a preliminary signal of potential desertification in HNP’s wetlands. First, the high variability of vegetation dynamics may hide underlying trends. Second, vegetation in arid and semiarid ecosystems is resilient to water limitations due to its spatial organization. In this case, the patch organization can serve as a preliminary metric of desertification, especially near tipping points (Barbier et al., 2006; Ge, 2023; Maestre and Escudero, 2009). The LMU approach gives essential insights into overall dynamics. However, it lacks spatial organization metrics that could enhance the interpretation of overall dynamics. Even if vegetation cover is stable, evaluating patch-size evolution could yield important insights into potential temporary WiD desertification processes in HNP.

A complex dynamic emerges at close range to the water’s edge. Here, our results show that bare soil exhibits a strong negative correlation with the water fraction. This negative correlation indicates that the closer you get to the water’s edge, the more the distribution of surface cover is influenced by water presence. In years of low water availability, such as during droughts, the surface cover vacated by receding water is primarily bare soil. This highlights the direct link between water presence and the exposure of bare soil in the inundation zone. Even though the statistical correlation between vegetation and water is highest at these short distances, vegetation does not entirely cover the area. This dynamic is particularly evident in our results for specific sites such as Roan Pan and White Hill, with Shapi and Togo showing similar trends, albeit to a lesser extent (Figure 10). These findings, at the pixel scale, show a highly dynamic proximal zone giving way to a more stable mosaic of vegetation and bare soil further away, aligning with earlier research on larger Southern African water bodies, which identified similar spatial distributions (Asare et al., 2018).

## 5. Conclusion

This study evaluated the evolution of water, vegetation, and bare soil around Wetlands in Drylands (WiDs) in Hwange National Park (HNP) from 1986 to 2021. These dynamics were analyzed using a linear multispectral unmixing (LMU) model based on Landsat 5, 7, and 8 imagery. The LMU model demonstrated a strong correlation with both the random forest classification and NDVI, lending confidence to the results. The analysis revealed a decline in water surface area in temporary wetlands between 1986 and 2021.

This decline was significantly and negatively correlated with temperature. This study provides strong evidence that a short hydroperiod is associated with increased vulnerability to climate change. In contrast, vegetation and bare soil did not show significant trends over the study period. However, the strong interdependence among these three land cover components underscores growing concerns as water becomes increasingly scarce in the park. Further in situ studies on vegetation patch size dynamics are needed to assess the potential risk of desertification in temporary wetlands across Southern African savannas, where water is a key factor in wetlands’ resilience to drought.

## Bibliography

Almalki, R., Khaki, M., Saco, P.M., Rodriguez, J.F., 2022. Monitoring and Mapping Vegetation Cover Changes in Arid and Semi-Arid Areas Using Remote Sensing Technology: A Review. Remote Sens. 14, 5143. 10.3390/rs14205143

Arraut, E.M., Loveridge, A.J., Chamaillé-Jammes, S., Valls-Fox, H., Macdonald, D.W., 2018. The 2013–2014 vegetation structure map of Hwange National Park, Zimbabwe, produced using free satellite images and software. KOEDOE - Afr. Prot. Area Conserv. Sci. 60. 10.4102/koedoe.v60i1.1497

Asare, F., Palamuleni, L.G., Ruhiiga, T., 2018. Land Use Change Assessment and Water Quality of Ephemeral Ponds for Irrigation in the North West Province, South Africa. Int. J. Environ. Res. Public. Health 15, 1175. 10.3390/ijerph15061175

Asner, G.P., Heidebrecht, K.B., 2002. Spectral unmixing of vegetation, soil and dry carbon cover in arid regions: Comparing multispectral and hyperspectral observations. Int. J. Remote Sens. 23, 3939–3958. 10.1080/01431160110115960

Asner, G.P., Lobell, D.B., 2000. A Biogeophysical Approach for Automated SWIR Unmixing of Soils and Vegetation. Remote Sens. Environ. 74, 99–112. 10.1016/S0034-4257(00)00126-7

Barbier, N., Couteron, P., Lejoly, J., Deblauwe, V., Lejeune, O., 2006. Self-organized vegetation patterning as a fingerprint of climate and human impact on semi-arid ecosystems. J. Ecol. 94, 537–547. 10.1111/j.1365-2745.2006.01126.x

Berg, A., McColl, K.A., 2021. No projected global drylands expansion under greenhouse warming. Nat. Clim. Change 11, 331–337. 10.1038/s41558-021-01007-8

Boix, D., Calhoun, A.J.K., Mushet, D.M., Bell, K.P., Fitzsimons, J.A., Isselin-Nondedeu, F., 2020. Conservation of Temporary Wetlands, in: Goldstein, M.I., DellaSala, D.A. (Eds.), Encyclopedia of the World’s Biomes. Elsevier, Oxford, pp. 279–294. 10.1016/B978-0-12-409548-9.12003-2

Ceballos, G., Ehrlich, P.R., Dirzo, R., 2017. Biological annihilation via the ongoing sixth mass extinction signaled by vertebrate population losses and declines. Proc. Natl. Acad. Sci. 114, E6089–E6096. 10.1073/pnas.1704949114

Chamaillé-Jammes, S., Fritz, H., Madzikanda, H., 2009. Piosphere contribution to landscape heterogeneity: a case study of remote-sensed woody cover in a high elephant density landscape. Ecography 32, 871–880. 10.1111/j.1600-0587.2009.05785.x

Chamaillé-Jammes, S., Fritz, H., Murindagomo, F., 2007a. Detecting climate changes of concern in highly variable environments: Quantile regressions reveal that droughts worsen in Hwange National Park, Zimbabwe. J. Arid Environ. 71, 321–326. 10.1016/j.jaridenv.2007.05.005

Chamaille-Jammes, S., Fritz, H., Murindagomo, F., 2006. Spatial patterns of the NDVI–rainfall relationship at the seasonal and interannual time scales in an African savanna. Int. J. Remote Sens. 27, 5185–5200. 10.1080/01431160600702392

Chamaillé-Jammes, S., Valeix, M., Fritz, H., 2007b. Managing heterogeneity in elephant distribution: interactions between elephant population density and surface-water availability: Surface water and elephant distribution. J. Appl. Ecol. 44, 625–633. 10.1111/j.1365-2664.2007.01300.x

Chang, M., Meng, X., Sun, W., Yang, G., Peng, J., 2021. Collaborative Coupled Hyperspectral Unmixing Based Subpixel Change Detection for Analyzing Coastal Wetlands. IEEE J. Sel. Top. Appl. Earth Obs. Remote Sens. 14, 8208–8224. 10.1109/JSTARS.2021.3104164

Cunnick, H., Ramage, J.M., Magness, D., Peters, S.C., 2023. Mapping Fractional Vegetation Coverage across Wetland Classes of Sub-Arctic Peatlands Using Combined Partial Least Squares Regression and Multiple Endmember Spectral Unmixing. Remote Sens. 15, 1440. 10.3390/rs15051440

Dzinotizei, Z., Murwira, A., Zengeya, F.M., Guerrini, L., 2018. Mapping waterholes and testing for aridity using a remote sensing water index in a southern African semi-arid wildlife area. Geocarto Int. 33, 1268–1280. 10.1080/10106049.2017.1343394

Feng, S., Fu, Q., 2013. Expansion of global drylands under a warming climate. Atmospheric Chem. Phys. 13, 10081–10094. 10.5194/acp-13-10081-2013

Furlan, L.M., Rosolen, V., Moreira, C.A., Bueno, G.T., Ferreira, M.E., 2021. The interactive pedological-hydrological processes and environmental sensitivity of a tropical isolated wetland in the Brazilian Cerrado. SN Appl. Sci. 3, 144. 10.1007/s42452-021-04174-7

Fynn, R.W.S., Murray-Hudson, M., Dhliwayo, M., Scholte, P., 2015. African wetlands and their seasonal use by wild and domestic herbivores. Wetl. Ecol. Manag. 23, 559–581. 10.1007/s11273-015-9430-6

Ge, Z., 2023. The hidden order of Turing patterns in arid and semi-arid vegetation ecosystems. Proc. Natl. Acad. Sci. 120, e2306514120. 10.1073/pnas.2306514120

Gorelick, N., Hancher, M., Dixon, M., Ilyushchenko, S., Thau, D., Moore, R., 2017. Google Earth Engine: Planetary-scale geospatial analysis for everyone. Remote Sens. Environ., Big Remotely Sensed Data: tools, applications and experiences 202, 18–27. 10.1016/j.rse.2017.06.031

Grenfell, S., Grenfell, M., Tooth, S., Mehl, A., O’Gorman, E., Ralph, T., Ellery, W., 2022. Wetlands in drylands: diverse perspectives for dynamic landscapes. Wetl. Ecol. Manag. 30, 607–622. 10.1007/s11273-022-09887-z

Guerschman, J.P., Scarth, P.F., McVicar, T.R., Renzullo, L.J., Malthus, T.J., Stewart, J.B., Rickards, J.E., Trevithick, R., 2015. Assessing the effects of site heterogeneity and soil properties when unmixing photosynthetic vegetation, non-photosynthetic vegetation and bare soil fractions from Landsat and MODIS data. Remote Sens. Environ. 161, 12–26. 10.1016/j.rse.2015.01.021

Gxokwe, S., Dube, T., Mazvimavi, D., 2020. Multispectral Remote Sensing of Wetlands in Semi-Arid and Arid Areas: A Review on Applications, Challenges and Possible Future Research Directions. Remote Sens. 12, 4190. 10.3390/rs12244190

Gxokwe, S., Mazvimavi, D., 2023. Available satellite data for monitoring small and seasonally flooded wetlands in semi-arid environments of southern Africa. Ecohydrology 17. 10.1002/eco.2605

Huang, H., Tuley, P.A., Tu, C., Zinnert, J.C., Rodriguez-Iturbe, I., D’Odorico, P., 2021. Microclimate feedbacks sustain power law clustering of encroaching coastal woody vegetation. Commun. Biol. 4, 1–7. 10.1038/s42003-021-02274-z

Huang, J., Ji, M., Xie, Y., Wang, S., He, Y., Ran, J., 2016a. Global semi-arid climate change over last 60 years. Clim. Dyn. 46, 1131–1150. 10.1007/s00382-015-2636-8

Huang, J., Yu, H., Guan, X., Wang, G., Guo, R., 2016b. Accelerated dryland expansion under climate change. Nat. Clim. Change 6, 166–171. 10.1038/nclimate2837

Hulot, F.D., Prijac, A., Lefebvre, J.-P., Msiteli-Shumba, S., Kativu, S., 2019. A first assessment of megaherbivore subsidies in artificial waterholes in Hwange National Park, Zimbabwe. Hydrobiologia 837, 161–175. 10.1007/s10750-019-3968-x

IPCC, 2022. Climate change 2022: Impacts, adaptation and vulnerability. Summary for policymakers.

Lange, R.T., 1969. The Piosphere: Sheep Track and Dung Patterns. 22. 10.2307/3895849

Maestre, F.T., Escudero, A., 2009. Is the patch size distribution of vegetation a suitable indicator of desertification processes? Ecology 90, 1729–1735. 10.1890/08-2096.1

Middleton, N., Thomas, D., UNEP, 1997. World atlas of desertification. Arnold.

Mitsch, W.J., Gosselink, J.G., 2007. Wetlands, Fourth Edition, John Wiley&Sons Inc. ed.

Mpakairi, K.S., 2019. Waterhole distribution and the piosphere effect in heterogeneous landscapes: evidence from north-western Zimbabwe. Trans. R. Soc. South Afr. 74, 219–222. 10.1080/0035919X.2019.1622607

Muñoz-Sabater, J., Dutra, E., Agustí-Panareda, A., Albergel, C., Arduini, G., Balsamo, G., Boussetta, S., Choulga, M., Harrigan, S., Hersbach, H., Martens, B., Miralles, D.G., Piles, M., Rodríguez-Fernández, N.J., Zsoter, E., Buontempo, C., Thépaut, J.-N., 2021. ERA5-Land: a state-of-the-art global reanalysis dataset for land applications. Earth Syst. Sci. Data 13, 4349–4383. 10.5194/essd-13-4349-2021

Nicholson, S.E., Farrar, T.J., 1994. The influence of soil type on the relationships between NDVI, rainfall, and soil moisture in semiarid Botswana. I. NDVI response to rainfall. Remote Sens. Environ. 50, 107–120. 10.1016/0034-4257(94)90038-8

Ozer, E., Leloglu, U.M., 2022. Wetland spectral unmixing using multispectral satellite images. Geocarto Int. 37, 15754–15777. 10.1080/10106049.2022.2102225

Parra, G., Guerrero, F., Armengol, J., Brendonck, L., Brucet, S., Finlayson, C.M., Gomes-Barbosa, L., Grillas, P., Jeppesen, E., Ortega, F., Vega, R., Zohary, T., 2021. The future of temporary wetlands in drylands under global change. Inland Waters 11, 445–456. 10.1080/20442041.2021.1936865

Prăvălie, R., 2016. Drylands extent and environmental issues. A global approach. Earth-Sci. Rev. 161, 259–278. 10.1016/j.earscirev.2016.08.003

Rasti, B., Zouaoui, A., Mairal, J., Chanussot, J., 2024. Image Processing and Machine Learning for Hyperspectral Unmixing: An Overview and the HySUPP Python Package. IEEE Trans. Geosci. Remote Sens. 62, 1–31. 10.1109/TGRS.2024.3393570

Rolls, R.J., Heino, J., Ryder, D.S., Chessman, B.C., Growns, I.O., Thompson, R.M., Gido, K.B., 2018. Scaling biodiversity responses to hydrological regimes. Biol. Rev. 93, 971–995. 10.1111/brv.12381

Roy, A., 2025. Structuring Influence of Wetlands in Drylands : From Water Availability Dynamics to Landscape Patterns in a Semi-Arid Southern African Savanna (Theses). Université Paris-Saclay.

Roy, A., Hulot, F.D., Soudani, K., 2025. Landsat-based remote sensing of surface water dynamics in southern African wetlands in drylands from 1986 to 2022. Ecosphere 16, e70332. 10.1002/ecs2.70332

Sandi, S.G., Rodriguez, J.F., Saintilan, N., Wen, L., Kuczera, G., Riccardi, G., Saco, P.M., 2020. Resilience to drought of dryland wetlands threatened by climate change. Sci. Rep. 10, 13232. 10.1038/s41598-020-70087-x

Schaffer-Smith, D., Swift, M., Killea, A., Brennan, A., Naidoo, R., Swenson, J.J., 2022. Tracking a blue wave of ephemeral water across arid southern Africa. Environ. Res. Lett. 17, 114063. 10.1088/1748-9326/ac98d9

Schmidt, H., Karnieli, A., 2000. Remote sensing of the seasonal variability of vegetation in a semi-arid environment. J. Arid Environ. 45, 43–59. 10.1006/jare.1999.0607

Sievers, M., Hale, R., Parris, K.M., Swearer, S.E., 2018. Impacts of human-induced environmental change in wetlands on aquatic animals. Biol. Rev. 93, 529–554. 10.1111/brv.12358

Slagter, B., Tsendbazar, N.-E., Vollrath, A., Reiche, J., 2020. Mapping wetland characteristics using temporally dense Sentinel-1 and Sentinel-2 data: A case study in the St. Lucia wetlands, South Africa. Int. J. Appl. Earth Obs. Geoinformation 86, 102009. 10.1016/j.jag.2019.102009

Thrash Derry, J.F., 1999. The nature and modelling of piospheres: a review. Koedoe 42, 73–94. 10.4102/koedoe.v42i2.234

Tooth, S., McCarthy, T.S., 2007. Wetlands in drylands: geomorphological and sedimentological characteristics, with emphasis on examples from southern Africa. Prog. Phys. Geogr. Earth Environ. 31, 3–41. 10.1177/0309133307073879

Valeix, M., 2011. Temporal dynamics of dry-season water-hole use by large African herbivores in two years of contrasting rainfall in Hwange National Park, Zimbabwe. J. Trop. Ecol. 27, 163–170. 10.1017/S0266467410000647

Williams, W. d., 1999. Conservation of wetlands in drylands: a key global issue. Aquat. Conserv. Mar. Freshw. Ecosyst. 9, 517–522. 10.1002/(SICI)1099-0755(199911/12)9:6%253C517::AID-AQC383%253E3.0.CO;2-C

Wu, X., Zhao, H., Wang, M., Yuan, Q., Chen, Z., Jiang, S., Deng, W., 2024. Evolution of Wetland Patterns and Key Driving Forces in China’s Drylands. Remote Sens. 16, 702. 10.3390/rs16040702

Xi, Y., Peng, S., Ciais, P., Chen, Y., 2021. Future impacts of climate change on inland Ramsar wetlands. Nat. Clim. Change 11, 45–51. 10.1038/s41558-020-00942-2

Xu, H., 2006. Modification of normalised difference water index (NDWI) to enhance open water features in remotely sensed imagery. Int. J. Remote Sens. 27, 3025–3033. 10.1080/01431160600589179

Xue, Z., Wang, Y., Huang, R., Yao, L., 2024. Study on Wetland Evolution and Landscape Pattern Changes in the Shaanxi Section of the Loess Plateau in the Past 40 Years. Land 13, 1268. 10.3390/land13081268

Yang, S., Yuan, Z., Ye, B., Zhu, F., Tang, X., Gao, R., Chu, Z., Liu, X., 2025. Niche partitioning and trait tradeoff strategies enable plants to coexist under interspecific competition in restored wetlands. Front. Plant Sci. 16. 10.3389/fpls.2025.1539136

Zedler, J.B., Kercher, S., 2005. WETLAND RESOURCES: Status, Trends, Ecosystem Services, and Restorability. Annu. Rev. Environ. Resour. 30, 39–74. 10.1146/annurev.energy.30.050504.144248

Zhang, X., Evans, J.P., Burrell, A.L., 2024. Less than 4% of dryland areas are projected to desertify despite increased aridity under climate change. Commun. Earth Environ. 5, 1–9. 10.1038/s43247-024-01463-y

Zhang, X., Liu, Y., Zhao, W., Li, J., Xie, S., Zhang, C., He, X., Yan, D., Wang, M., 2023. Impact of Hydrological Changes on Wetland Landscape Dynamics and Implications for Ecohydrological Restoration in Honghe National Nature Reserve, Northeast China. Water 15, 3350. 10.3390/w15193350

Zhao, F., Ma, S., Wu, Y., 2021. Changes in Dry-Season Water Availability and Attributions in the Yellow River Basin, China. Front. Environ. Sci. 9. 10.3389/fenvs.2021.762137

Zhou, S., Williams, A.P., Lintner, B.R., Berg, A.M., Zhang, Y., Keenan, T.F., Cook, B.I., Hagemann, S., Seneviratne, S.I., Gentine, P., 2021. Soil moisture–atmosphere feedbacks mitigate declining water availability in drylands. Nat. Clim. Change 11, 38–44. 10.1038/s41558-020-00945-z

