## Supplementary Materials for "Hydroperiod buffers water surface decline in dryland wetlands: A 36-year analysis in Hwange National Park"

**Table S1**: Validation confusion matrix of the Sentinel-2 Random Forest classification

| Reference | FreeWaterBodies | Wetlands | Herbaceous Crops | Shrubs Trees | BareSoils |
|---|---|---|---|---|---|
| FreeWaterBodies | 221 | 0 | 0 | 0 | 0 |
| Wetlands | 0 | 2385 | 0 | 0 | 0 |
| Herbaceous Crops | 0 | 0 | 325 | 5 | 0 |
| Shrubs Trees | 0 | 1 | 18 | 2393 | 0 |
| BareSoils | 0 | 0 | 1 | 1 | 185 |

**Table S2**: Water fraction autocorrelation depending on the distance or the hydroperiod.

| sType | Label | Lag | ACF | p-value |
|---|---|---|---|---|
| Distance | 50 m | 1 | -0.11167576026209888 | 0.4906883555653354 |
| Distance | 50 m | 2 | -0.16950434699576783 | 0.44876339516408725 |
| Distance | 50 m | 3 | 0.12563586769846985 | 0.5238608472534576 |
| Distance | 50 m | 4 | -0.06278669396723693 | 0.661547992957779 |
| Distance | 100 m | 1 | -0.016144155764163656 | 0.9206343174313847 |
| Distance | 100 m | 2 | -0.01821699413779599 | 0.9885905733786945 |
| Distance | 100 m | 3 | 0.021448641688098336 | 0.9977739268204653 |

| | | | | |
|---|---|---|---|---|
| Distance | 100 m | 4 | -0.14764808247895675 | 0.9169489622084961 |
| Distance | 150 m | 1 | 0.031261097581413905 | 0.84701383033222 |
| Distance | 150 m | 2 | 0.07183562679060584 | 0.8870415564516471 |
| Distance | 150 m | 3 | -0.006340053848351511 | 0.970652474631397 |
| Distance | 150 m | 4 | -0.1641850945197929 | 0.8498321877461599 |
| Distance | 200 m | 1 | 0.05099923470962204 | 0.752955139411077 |
| Distance | 200 m | 2 | 0.11775829279094269 | 0.724973688535633 |
| Distance | 200 m | 3 | -0.0114292927873219 | 0.8852389617277967 |
| Distance | 200 m | 4 | -0.16906726403613373 | 0.7646819398741438 |
| Distance | 250 m | 1 | 0.05880296363455956 | 0.7166747974256069 |
| Distance | 250 m | 2 | 0.14840322904886158 | 0.6077579057561135 |
| Distance | 250 m | 3 | -0.016713974089572068 | 0.7994946422843141 |
| Distance | 250 m | 4 | -0.16852943965969683 | 0.7001754441793042 |
| Distance | 300 m | 1 | 0.0574889152808775 | 0.7227419202177104 |
| Distance | 300 m | 2 | 0.17425723003016566 | 0.5174978699757804 |
| Distance | 300 m | 3 | -0.03379898065213999 | 0.7140581855694753 |
| Distance | 300 m | 4 | -0.154676160336468 | 0.6692937908228019 |
| Distance | 350 m | 1 | 0.04685793017722137 | 0.7724384015207684 |
| Distance | 350 m | 2 | 0.18550093045220817 | 0.488218561240071 |
| Distance | 350 m | 3 | -0.04328640883918238 | 0.6800078069510065 |
| Distance | 350 m | 4 | -0.14249455382855028 | 0.6702260213020608 |
| Distance | 400 m | 1 | 0.04516003238061793 | 0.7804688065909147 |
| Distance | 400 m | 2 | 0.1910630753307644 | 0.46995731717166356 |
| Distance | 400 m | 3 | -0.03896208746725805 | 0.6658326215348798 |
| Distance | 400 m | 4 | -0.1452662818995467 | 0.6530367466926102 |

| | | | | |
|---|---|---|---|---|
| FreqClass | 0-10% | 1 | 0.215505256079379 | 0.18351660482174809 |
| FreqClass | 0-10% | 2 | 0.3308803696632824 | 0.04818908568503618 |
| FreqClass | 0-10% | 3 | -0.07375348586222008 | 0.09852199406023761 |
| FreqClass | 0-10% | 4 | -0.062457522983873984 | 0.1680748147179332 |
| FreqClass | 10-30% | 1 | -0.003172450598929547 | 0.9843792238433893 |
| FreqClass | 10-30% | 2 | 0.07575324872248633 | 0.8933390746545322 |
| FreqClass | 10-30% | 3 | 0.02118041790173037 | 0.9702384042977874 |
| FreqClass | 10-30% | 4 | -0.20785142305199278 | 0.726844098652067 |
| FreqClass | 30-50% | 1 | 0.05602454005381246 | 0.7295237012591187 |
| FreqClass | 30-50% | 2 | -0.025853578509492423 | 0.9297033960623896 |
| FreqClass | 30-50% | 3 | 0.05842178559233258 | 0.9630268249133275 |
| FreqClass | 30-50% | 4 | -0.22380364512405176 | 0.6669158158865488 |
| FreqClass | 50-70% | 1 | -0.03673137908947579 | 0.8206646827673276 |
| FreqClass | 50-70% | 2 | 0.006532163169947449 | 0.9738174325573551 |
| FreqClass | 50-70% | 3 | 0.026951851102687045 | 0.9938558562130041 |
| FreqClass | 50-70% | 4 | -0.05175631223062913 | 0.9955730506717361 |
| FreqClass | 70-100% | 1 | 0.0638034222134215 | 0.693753015170683 |
| FreqClass | 70-100% | 2 | 0.07620516600620994 | 0.8257434939194385 |
| FreqClass | 70-100% | 3 | -0.12752065051122555 | 0.7913263692469333 |
| FreqClass | 70-100% | 4 | -0.09945980160486041 | 0.8347105584923944 |

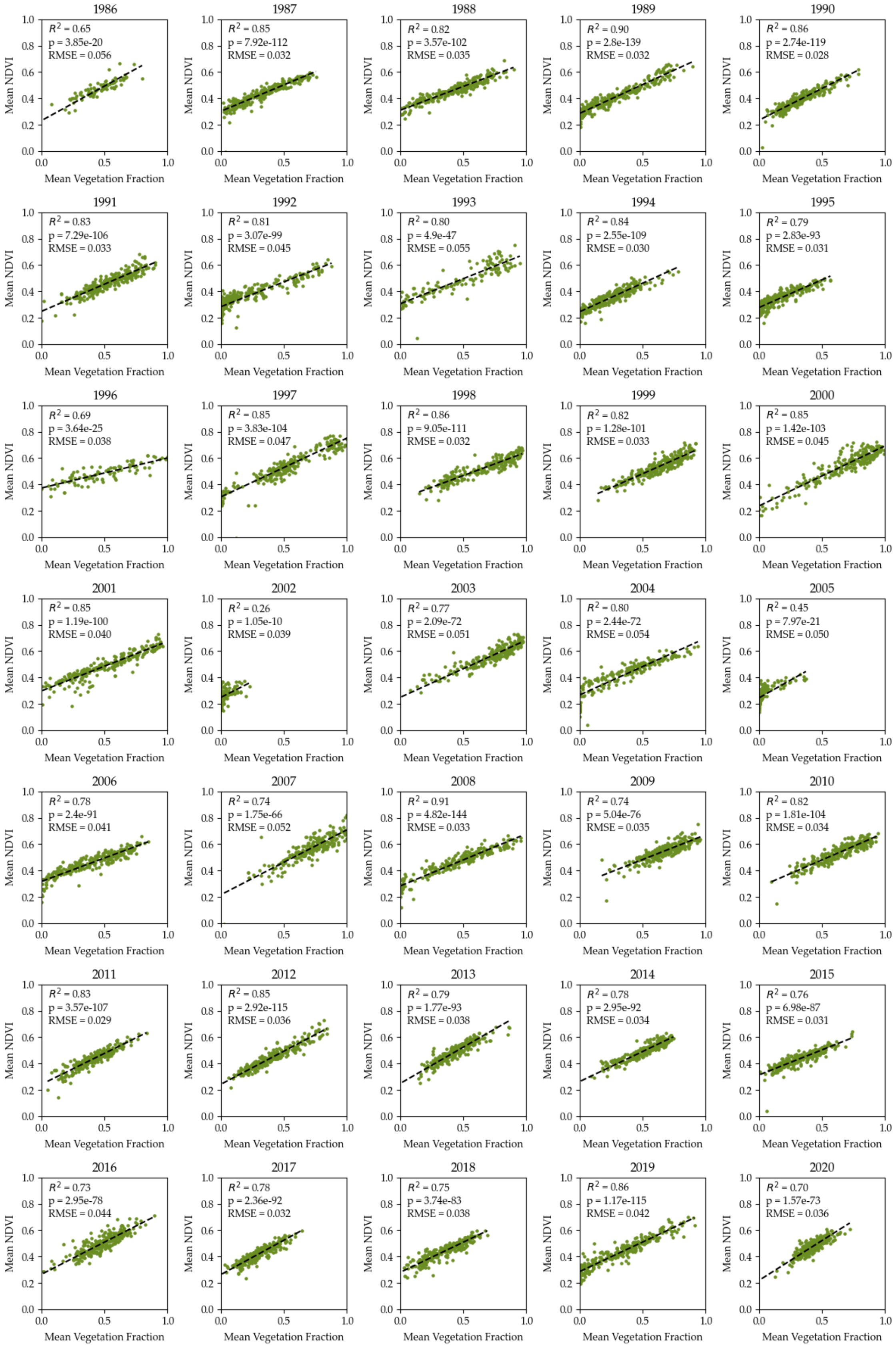


**Figure S1:** Linear regression between the vegetation fraction extracted from the Linear Multispectral Unmixing and the NDVI from 1986 to 2020. $R^2$, p-value, and RMSE are mentioned on the plot.

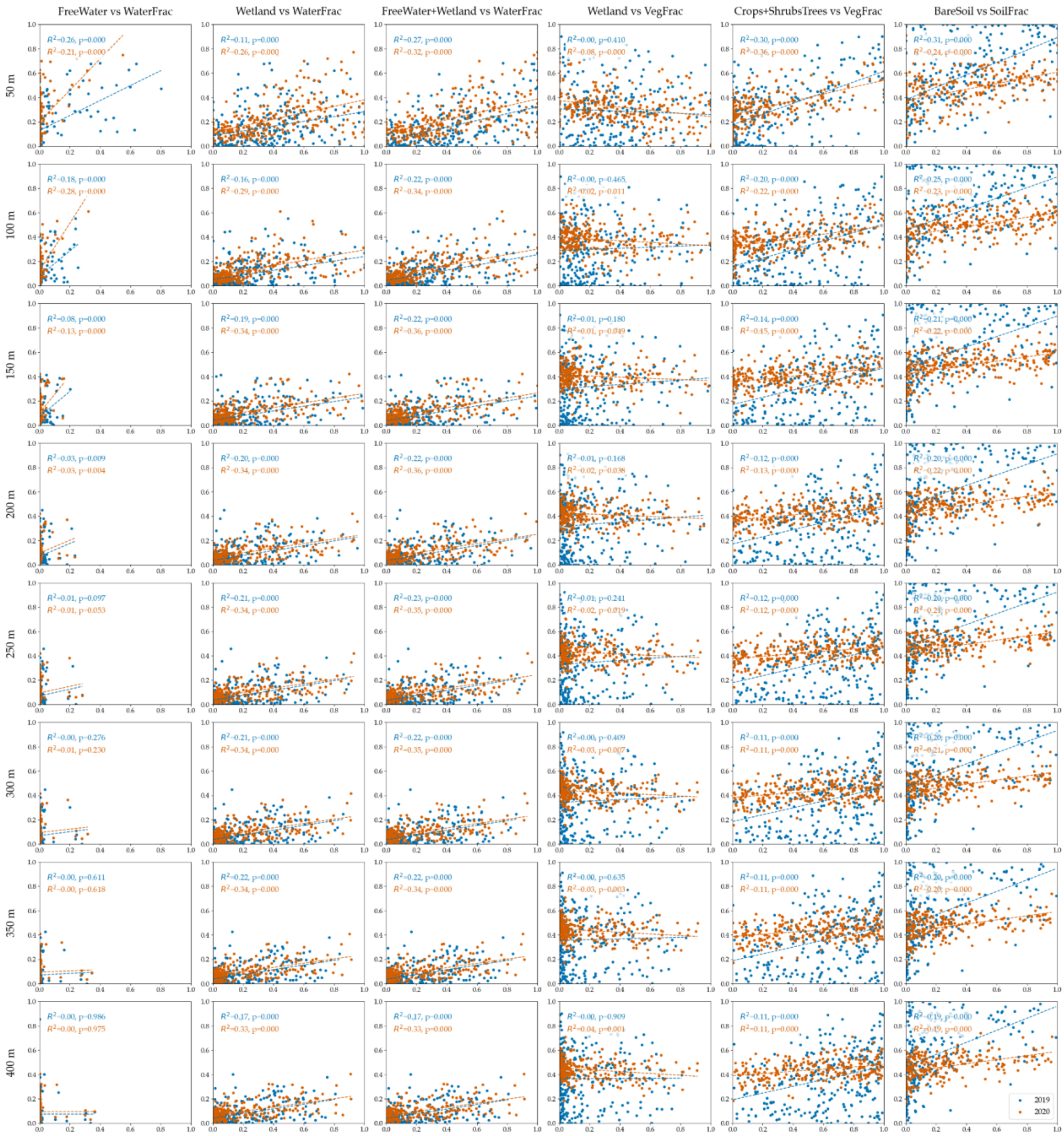


**Figure S2:** Correlation between each fraction and each Random Forest classification pixel count, from 50m to 400m. The RF pixel count is on the X axis and the LMU fraction is on the Y axis. Each comparison is specified above the column. 2019 is in blue and 2020 is in orange.
